# Substrate-induced modulation of protein-protein interactions within human mitochondrial cytochrome P450-dependent system

**DOI:** 10.1101/2020.11.13.381095

**Authors:** E.O. Yablokov, T.A. Sushko, L.A. Kaluzhskiy, A.A Kavaleuski, Y.V. Mezentsev, P.V. Ershov, A.A Gilep, A.S. Ivanov, N.V. Strushkevich

## Abstract

Steroidogenesis is strictly regulated at multiple levels, as produced steroid hormones are crucial to maintain physiological functions. Cytochrome P450 enzymes are key players in adrenal steroid hormone biosynthesis and function within short redox-chains in mitochondria and endoplasmic reticulum. However, mechanisms regulating supply of reducing equivalents in the mitochondrial CYP-dependent system are not fully understood. In the present work, we aimed to estimate how the specific steroids, substrates, intermediates and products of multistep reactions modulate protein-protein interactions between adrenodoxin (Adx) and mitochondrial CYP11s. Using the SPR technology we determined that steroid substrates affect affinity and stability of CYP11s – Adx complexes in an isoform-specific mode. In particular, cholesterol induces a 4-fold increase in the rate of CYP11A1 – Adx complex formation without significant effect on dissociation (k_off_ decreased ~1.5-fold), overall increasing complex affinity. At the same time steroid substrates decrease the affinity of both CYP11B1 – Adx and CYP11B2 – Adx complexes, predominantly reducing their stability (4-7 fold). This finding reveals differentiation of protein-protein interactions within the mitochondrial pool of CYPs, which have the same electron donor. The regulation of electron supply by the substrates might affect the overall steroid hormones production. Our experimental data provide further insight into protein-protein interactions within CYP-dependent redox chains involved in steroidogenesis.

## 1. Introduction

Steroid hormones control important physiological functions in humans. Steroidogenesis is tightly regulated at multiple levels, including transcriptional, translational, and post-translational [1–3]. Proteinprotein interactions (PPI), involved in electron transfer, modulate catalytic activity of steroidogenic enzymes, while substrate availability determines the level of produced hormones [4,5]. Compartmentalization of the components of the steroidogenic machinery is also important for the biosynthesis of different classes of hormones [6]. The mitochondria of the adrenal steroidogenic cells contains short redox chains, comprising of cytochrome P450 enzymes (CYP) CYP11A1, CYP11B1, CYP11B2, and their cognate electron-transfer partners, adrenodoxin reductase (AdR) and Adx. CYP11A1 catalyses cholesterol side chain cleavage yielding pregnenolone, the precursor of all steroid hormones. CYP11B1 and CYP11B2 produce gluco-and mineralocorticoids, respectively, regulating homeostasis, stress response, salt and water balance, growth. That is why their catalytic activity is strictly regulated [1].

Electron transfer is crucial step in the CYP catalytic cycle, hence the interaction of CYP with redox-partner is required for the catalytic activity [5]. In many CYPs ligand binding induces structural rearrangements, [7–11] which in turn affect the redox partner binding [12]. The effect of substrates was previously reported for the PPI in microsomal CYP-dependent systems [13–16]. For the mitochondrial steroidogenic CYPs this effect is less studied. Reports about CYP24A1, involved in calcitriol metabolism, revealed that vitamin D analogue induced structural changes in this mitochondrial CYP, decreasing its binding affinity to Adx [12,17]. Since the oxidation of physiologically significant substrates is a multistep process for many CYPs, it is of special interest to explore the regulation of the individual reactions. CYP11A1 was extensively studied for decades and considered as a model for the multistep reactions and for the electron transfer in P450-dependent systems [7–9,11]. The impact of redox partner binding on the interaction of CYP11A1 with substrates and intermediates was addressed previously [18,19]. Significantly less is known about CYP11B isoforms, which recently gained much interest as drug targets. Moreover, it is difficult to compare results obtained using different experimental conditions and systems. The estimated Adx content in adrenals might not be sufficient to supply electrons for all steroidogenic enzymes [20,21], suggesting some regulatory mechanisms [5]. Here, we applied surface plasmon resonance (SPR) analysis to study the effect of not only the substrates, but also intermediates of multistep reactions, as well as the products on the PPI between mitochondrial steroidogenic CYPs and Adx. Our data reveal different effects of steroids on affinity and stability of Adx complexes with three studied CYPs. Substrate-induced changes in PPI might contribute to the regulation of steroid hormones production.

## 2. Materials and methods

### 2.1 Equipment

Real-time measurements of PPIs were performed on Biacore T200 and Biacore 8K SPR biosensors (GE Healthcare, USA) using CM4 optical chips coated with a layer of carboxymethylated dextran. The sensorgrams of PPIs were registered as a continuous record of the biosensor signal in resonance units, RU. 1 RU is approximately equal to the binding of 1 pg of protein per 1 mm^2^ of the chip surface. Proteins purification procedures were performed using the chromatographic system Akta Purifier (GE Healthcare, USA).

### 2.2 Chemicals

The following reagents were used: progesterone, pregnenolone, 11-deoxycorticosterone, 11-deoxycortisol, cortisol, cholesterol, 22R-hydroxycholesterol, 20R,22R-dihydroxycholesterol, 20S-hydroxycholesterol, 25-hydroxycholesterol, CHAPS, NADPH, HEPES, TRIS, DLPC, EDTA (Sigma-Aldrich, USA); Ni-NTA-Agarose (Qiagen, USA); Emulgen 913 (Kao Atlas, Japan); dithiothreitol (Gibco BRL, USA). Special reagents for biosensor measurements were obtained from GE Healthcare (GE Healthcare, USA): HBS-EP+ buffer (150 mM NaCl, 10 mM HEPES, 3 mM EDTA, 0.05% P-20 surfactant, pH 7.4), HBS-N buffer (150 mM NaCl, 10 mM HEPES, pH 7.4), 50 mM NaOH solution, 70% glycerol solution, 10 mM acetate buffer (pH 4.5), the amine-coupling kit for covalent immobilization of proteins on the CM4 optical chip: 1-ethyl-3-(3-dimethylaminopropyl) carbodiimide-HCl (EDC), N-hydroxysuccinimide (NHS), 1M ethanolamine-HCl (pH 8.5). Other reagents of analytical purity were obtained from local suppliers.

### 2.3 Proteins

In the current study, recombinant highly purified (> 95% by SDS-PAGE) mitochondrial human CYP11A1, CYP11B1 and CYP11B2 and their redox partners Adx and AdR were used. CYP11A1 and CYP11B isoforms were expressed and purified as described previously [22–24]. Recombinant Adx and AdR were purified according to [23]. Spectral studies confirmed that CYP11A1, CYP11B1, CYP11B2 were low-spin and ligand-free, containing P450 form (>75%) according to CO-difference spectra [25]. Enzymatic activity of CYPs were measured in the reconstituted system as described previously [22,24] using cholesterol as a substrate for CYP11A1, 11-deoxycorticosterone for CYP11B2 and 11-deoxycortisol for CYP11B1.

### 2.4 Immobilization of Adx on the CM4 optical chip

Adx was covalently immobilized on the surface of the CM4 chip according to the standard amino-coupling protocol. HBS-N (150 mM NaCl, 10 mM HEPES, pH 7.4) was used as a running buffer. The carboxyl groups of dextran were activated by injecting a mixture of 0.4 M EDC and 0.1 M NHS for 7 min at a flow rate of 5 μl/min. The protein sample diluted to 15 μg/ml in 10 mM sodium acetate (pH 4.5) was injected for 10 min at a flow rate of 2 μl/min. Finally, ethanolamine-HCl (1.0 M) solution was injected for 1 min at a flow rate of 5 μl/min to block the unreacted carboxyl groups of dextran. The amount of Adx immobilized on the chip surface was approximately 150 fmol.

### 2.5 SPR analysis of PPIs between CYPs and immobilized Adx

The interactions of CYP isoforms with immobilized Adx were measured in a range from 25 to 250 nM and at 25°C. The analyzed CYP was injected into the control (without immobilized proteins) and working channels of the optical biosensor for 7 min at a flow rate of 10 μl/min. HBS-EP+, containing 0.5% ethanol was used as a running buffer.

The resulting sensorgrams were obtained by subtracting the signals of the control channel and the working channel. Regeneration of the chip surface was performed after each protein injection using a 10 mM HEPES (pH 7.4) buffer, containing 1 M NaCl and 0.2% CHAPS for 1 min at a flow rate of 50 μl/min. The data were fitted to a Langmuir 1:1 global fitting model using Biacore BIAevaluation software v. 4.1 (GE Healthcare, USA) to calculate the association and dissociation rate constants (k_on_ and k_off_, respectively) and the apparent equilibrium dissociation constant (K_d_). K_d_ was calculated as K_d_ = k_off_/k_on_. Significance of differences between the mean values of τ_1/2_, kinetic and equilibrium constants obtained in the presence and absence of substrates was determined by Student’s t-test. Steroid compounds dissolved in ethanol were added to the sample to the final concentration of 100 μM (the final concentration of ethanol in the experiments was not higher than 0.5%), used for biochemical assays and ensures fully occupied steroid-binding site of the enzymes [22,24]. To obtain sensorgrams of Adx – CYP interactions without steroids, equal amounts of ethanol were added to the samples. Before injection all samples were incubated for 30 min at 25°C. All measurements were performed in triple replicates. The SPR analysis showed that the background binding levels of compounds (100 μM) to the immobilized Adx did not exceed 2 – 5 RU, indicating the absence of any interaction between steroid compounds and Adx. τ_1/2_ values, characterizing half-time dissociation of a protein-protein complex, were calculated from k_off_ values according to the equation (1).

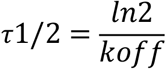

## 3. Results

### 3.1 Catalytic activity of recombinant proteins

The catalytic activity of CYP11A1, CYP11B1, CYP11B2 was measured in the reconstituted system with the reaction rates (turnover number (**TN**)) for CYP11A1 equal to 10.7 ± 1.3 min^-1^, CYP11B1 TN=96.8 ± 9.3 min^-1^, and CYP11B2 TN=37.6 ± 2.4 min^-1^, which are comparable with previously reported values [22,24]. The activity was measured to ensure that the enzymes used for designed SPR studies were catalytically competent.

### 3.2 Characterization of CYP11 – Adx interactions by SPR

Effective application of SPR biosensors for the CYP-containing systems has been reported earlier [13,26–29]. Here we aimed to dissect it to more details, specifically, to investigate how different steroids contribute to the PPI between the three mitochondrial CYP11s: CYP11A1 – cholesterol side chain cleavage enzyme, CYP11B1 – 11ß-hydroxylase and CYP11B2 – aldosterone synthase and their common redox partner Adx.

#### 3.2.1 Ligand-mediated interactions of CYP11A1 with Adx

Human CYP11A1 converts cholesterol to pregnenolone, which serves as the metabolic precursor for all adrenal steroids. The reaction occurs in three steps. The first two steps are stereospecific hydroxylations leading to the formation of 22R-hydroxycholesterol and 20R,22R-dihydroxycholesterol, respectively, followed by the oxidative scission of the C20–22 bond and pregnenolone formation (Fig. 1) [22]. We compared the effect of cholesterol substrate and intermediates (22R-hydroxycholesterol and 20R,22R-dihydroxycholesterol) on the affinity of CYP11A1 – Adx complex. Typical sensorgrams of CYP – Adx interactions as well as sensorgrams in the presence of steroid compounds are presented on Fig. 2. Calculated K_d_, k_on_, k_off_, and the half-life time of complex dissociation (τ_1/2_) values are summarized in the Table 1. Twofold change of a parameter value was considered to be significant (p <0.01). Cholesterol decreased the resulting K_d_ value of CYP11A1 – Adx complex predominantly due to a 4-fold increase in k_on_, while k_off_ changed ~1.5-fold (Table 1, Fig. 3). This is in contrast with previously published data [11], where cholesterol did not increase affinity of CYP11A1 to Adx. It could be explained by different experimental settings. More specifically, there were differences in the concentrations of both steroids and protein analytes, in the preincubation time with steroids as well as the level of adrenodoxin immobilization on the chip. The first intermediate (22R-hydroxycholesterol) did not change the parameters of the interaction of CYP11A1 – Adx. In the presence of the second intermediate (20R,22R-dihydroxycholesterol) K_d_ value of the CYP11A1– Adx complex increased about 3.4 times due to 2-fold decrease in the k_on_ and about 1.8-fold increase in k_off_. Compared to cholesterol, reaction intermediates 22R-hydroxycholesterol and 20R,22R-dihydroxycholesterol increased the K_d_ of the CYP11A1 – Adx complex 4 times and almost 10-fold, respectively. A decrease in the k_on_ of the CYP11A1 – Adx complex was observed in the following sequence cholesterol > 22R-hydroxycholesterol > 20R,22R-dihydroxycholesterol, indicating that binding of the intermediates decreased the rate of the complex formation (Fig. 1, Fig 3.1).

**Fig. 1.**
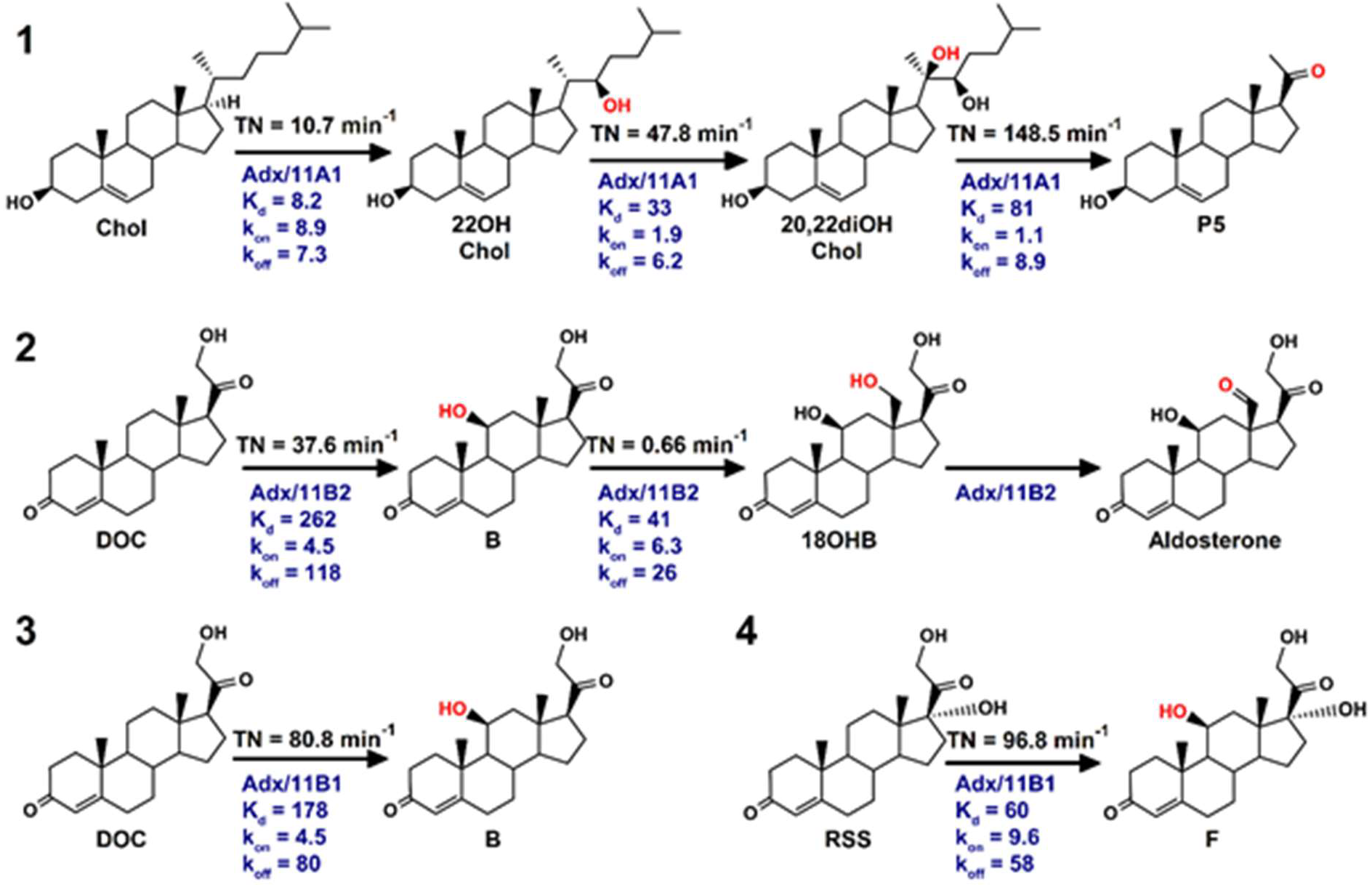
Catalytic activity of mitochondrial steroidogenic CYP11s. Cleavage of the cholesterol side chain by CYP11A1 (1), synthesis of aldosterone by CYP11B2 (2), enzymatic activity of CYP11B1 (3,4). Catalytic activity (TN, min^-1^) of human CYP11A1 [22] and CYP11B [24], is shown above arrows. Rate constants k_on_ (10^4^ M^-1^s^-1^), k_off_ (10^-4^ s^-1^) and apparent equilibrium constant (K_d_, nM) of CYP – Adx interaction in the presence of appropriate steroids are indicated below arrows. Chol – cholesterol, 22OHChol – 22R-hydroxycholesterol, 20,22diOHChol – 20R,22R-dihydroxycholesterol, P5 – pregnenolone, DOC – 11-deoxycorticosterone, B – corticosterone, 18OHB – 18-hydroxycorticosterone, RSS — 11-deoxycortisol, F – cortisol. **Double column**

**Fig. 2.**
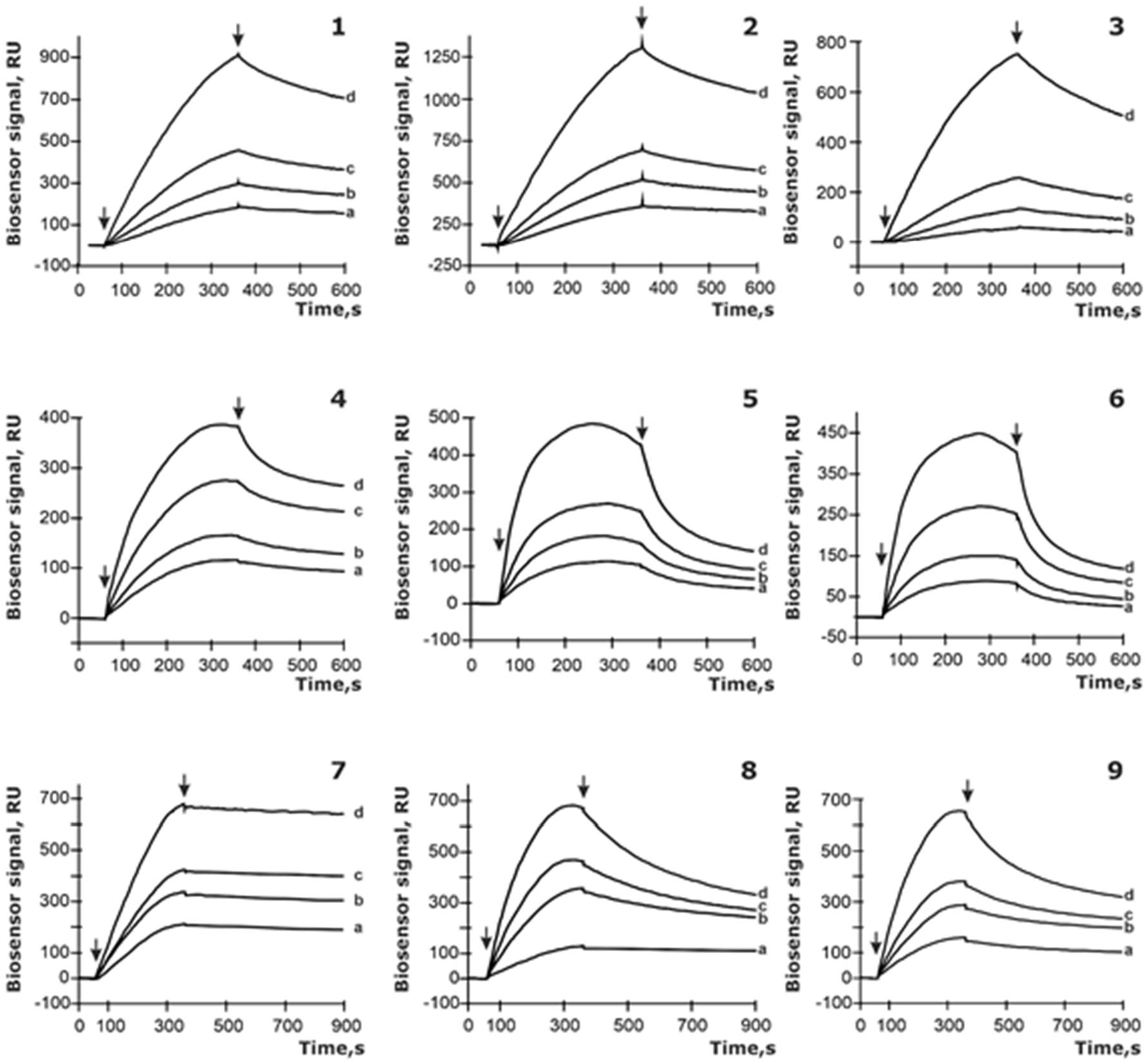
Sensorgrams of interaction CYP – Adx at 25°C in the HBS-EP+ buffer. CYP’s concentration: a – 25 nM, b – 50 nM, c – 125 nM, d – 250 nM. 1 – CYP11A1 – Adx; 2 – CYP11A1 – Adx in the presence of 100 μM cholesterol; 3 – CYP11A1 – Adx in the presence of 100 μM pregnenolone; 4 – CYP11B1 – Adx; 5 – CYP11B1 – Adx in the presence of 100 μM 11-deoxycorticosterone; 6 – CYP11B1 – Adx in the presence of 100 μM 11-deoxycortisol; 7 – CYP11B2 – Adx; 8 – CYP11B2 – Adx in the presence of 100 μM 11-deoxycorticosterone; 9 – CYP11B2 – Adx in the presence of 100 μM 11-deoxycortisol. Arrows indicate the start and the end point of CYP sample’s injections. **Double column**

**Table 1.** The effect of steroids on CYP11A1 – Adx interaction: complex affinity (apparent K_d_), association and dissociation rate constants (k_on_, k_off_), dissociation half-time (τ_1/2_).

| Compound | $K_d$<br>(n M) | $k_{on}$<br>($10^4 \text{ M}^{-1} \text{ s}^{-1}$) | $k_{off}$<br>($10^{-4} \text{ s}^{-1}$) | $\tau_{1/2}$<br>(s) |
| --- | --- | --- | --- | --- |
| Ligand-free | 24±2 | 2.1±0.3 | 5.0±0.6 | 1380±210 |
| Chol (S) | 8.2±1.3<br>↓* | 8.9±1.1 ↑* | 7.3±1.1 | 945±138 |
| 22OHChol (I) | 33±4 | 1.9±0.2 | 6.2±0.8 | 1113±164 |
| 20,22diOHChol (I) | 81±5 ↑* | 1.1±0.1 ↓* | 8.9±1.2 * | 775±108* |
| 25OHChol (S) | 110±10<br>↑* | 1.0±0.1 ↓* | 11±2 ↑* | 627±92 ↓* |
| 20OHChol (S) | 75±7 ↑* | 1.1±0.1 ↓* | 8.3±1.1 * | 831±119* |
| DOC (C) | 23±3 | 2.8±0.2 | 6.3±0.8 | 1095±155 |
| RSS (C) | 28±4 | 2.6±0.2 | 7.3±1.2 | 945±138 |
| P5 (P) | 172±21<br>↑* | 0.50±0.06 ↓* | 8.6±1.3* | 802±116 |
| P4 (C) | 20±3 | 3.3±0.4 | 6.5±0.7 | 1062±153 |

The direction of the significant changes of constants in the presence of compounds was shown with arrows. S – substrate, I – intermediate, P – product, C – negative control, steroid that does not bind to the active site of the CYP, «Ligand-free» – interaction of CYP in the ligand-free form. Chol – cholesterol, 22OHChol - 22R-hydroxycholesterol, 20,22diOHChol - 20R,22R-dihydroxycholesterol, 25OHChol - 25-hydroxycholesterol, 20OHChol - 20S-hydroxycholesterol, DOC – 11-deoxycorticosterone, RSS — 11-deoxycortisol, P4 – progesterone, P5 – pregnenolone.
Values of complex formation constants represented as mean ±SD (Standard deviation), (n = 9)
\* the value is significantly different from the corresponding value of the CYP– Adx interaction without steroid compounds, with $p < 0.01$

We also analyzed the effect of alternative substrates, 25-hydroxycholesterol and 20S-hydroxycholesterol, on CYP11A1 – Adx interaction. 25-hydroxycholesterol and 20S-hydroxycholesterol induced two-fold decrease in k_on_ compared to the ligand-free complex, with K_d_ increased 4.6 times and 3.1 times, respectively. However, the rate of the complex formation and the affinity of the complex decreased almost 10 times when compared with cholesterol, indicating the preferable interaction of complex CYP11A1– cholesterol with Adx.

In the presence of the final product (pregnenolone) we detected the inhibition of CYP11A1 – Adx complex formation. The K_d_ of CYP11A1–Adx complex increased almost 21-fold compared to the initial substrate cholesterol, and was mainly attributed to k_on_ value. The same tendency of pregnenolone effect, namely, decreasing k_on_ and increasing the K_d_ values of the CYP11A1 – Adx complex, although to a lesser extent, was observed compared to the ligand-free CYP11A1. The non-specific steroids, including substrates, intermediates, and products of CYP11B1 and CYP11B2 reactions which are not metabolized by CYP11A1, did not cause any effect on the CYP11A1 – Adx complex.

Thus, we detected distinct effects of the substrates, intermediates, and products on the complex CYP11A1 – Adx, mainly reflected in the association rate values, while the contribution of k_off_ was not significant.

#### 3.2.2. Ligand-mediated interactions of CYP11B enzymes with Adx

CYP11B1 converts 11-deoxycortisol to cortisol, while CYP11B2 catalyzes the biosynthesis of aldosterone. CYP11B2 has not only 11β-hydroxylase activity, but also 18-hydroxylase and 18-oxidase activities, producing aldosterone via three-step catalysis (Fig. 1) [24]. Both 11-deoxycortisol and 11-deoxycorticosterone are substrates for CYP11B1 and CYP11B2 *in vitro,* while anatomic zonation of adrenal cortex and different location of CYP11B1 and CYP11B2 prevent the access of 11-deoxycortisol to zona reticularis, where CYP11B2 is expressed. In our experimental setup we analyzed the effect of various steroids on the parameters of both CYP11B1 – Adx and CYP11B2 – Adx complex formation to estimate their contribution. The 11-deoxycortisol increased both association (4-fold) and dissociation (~ 3.2-fold) rates of CYP11B1 – Adx complex (Table 2). As a result, 11-deoxycortisol did not affect the affinity of the CYP11B1 – Adx complex, as the equal in amplitude, but opposite changes in k_on_ and k_off_ did not change the K_d_ value (Fig. 3.2). 11-deoxycorticosterone increased k_off_ of the CYP11B1 – Adx complex 4.4-fold and k_on_ increased almost 2-fold, so the resulting K_d_ increased 2.4 times. Reaction products corticosterone and cortisol as well as cholesterol did not affect significantly the interaction between CYP11B1 and Adx.

**Table 2.** The influence of the steroid compounds on CYP11B – Adx interaction parameters: complex affinity (apparent K_d_), association and dissociation rate constants (k_on_, k_off_), half-time of the complex dissociation (τ_1/2_).

| CYP | Compound | $K_d$<br>(nM) | $k_{\text{on}}$<br>($10^4 \text{ M}^{-1} \text{ s}^{-1}$) | $k_{\text{off}}$<br>($10^{-4} \text{ s}^{-1}$) | $\tau_{1/2}$<br>(s) |
| --- | --- | --- | --- | --- | --- |
| CYP11B1 | Ligand-free | 75±6 | 2.4±0.3 | 18±1 | 383±56 |
|  | DOC (S) | 178±10 ↑* | 4.5±0.5 ↑* | 80±10 ↑* | 86±12 ↓* |
|  | RSS (S) | 60±8 | 9.6±1.1 ↑* | 58±9 ↑* | 119±18 ↓* |
|  | Chol (C) | 88±12 | 2.6±0.2 | 23±3 | 300±43 |
|  | P4 (S) | 207±19 ↑* | 1.4±0.1 | 29±3 | 238±33 |
|  | B (C) | 54±8 | 2.8±0.2 | 15±1 | 460±68 |
|  | F (P) | 59±5 | 2.7±0.2 | 16±2 | 431±64 |
| CYP11B2 | Ligand-free | 24±1 | 7.0±0.6 | 17±2 | 406±60 |
|  | DOC (S) | 262±21 ↑* | 4.5±0.5 | 118±15 ↑* | 58±8 ↓* |
|  | RSS (S) | 113±8 ↑* | 7.2±0.9 | 81±12 ↑* | 85±11 ↓* |
|  | Chol (C) | 25±3 | 6.8±0.6 | 17±2 | 406±56 |
|  | P4 (S) | 28±2 | 18±2 ↑* | 50±7 ↑* | 138±18 ↓* |
|  | B (I) | 41±5 | 6.3±0.6 | 26±2 | 265±38 |
|  | F (P) | 32±6 | 7.3±0.9 | 23±2 | 300±39 |

The direction of the significant changes of constants in the presence of compounds was shown with arrows. S – substrate, I – intermediate, P – product, C – negative control, steroid that does not bind to the
active site of the CYP, «Ligand free» – interaction with CYP in the ligand-free form. Chol – cholesterol, DOC – 11-deoxycorticosterone, RSS – 11-deoxycortisol, B – corticosterone, F – cortisol, P4 – progesterone.
Values of complex formation constants represented as mean ± SD (Standard deviation), (n = 9)
\* the value is significantly different from the CYP – Adx interaction parameters obtained without steroid compounds, with $p < 0.01$

11-deoxycortisol increased the K_d_ of the CYP11B2 – Adx complex 4.7 times due to the proportional increase in the rate of the complex dissociation. Similarly, 11-deoxycorticosterone increased K_d_ of the CYP11B2 – Adx complex 11-fold due to the 7-fold increase in k_off_. Both 11-deoxycortisol and 11-deoxycorticosterone did not affect the CYP11B2 – Adx complex formation, k_on_ was increased but only slightly. Intermediate corticosterone did not significantly (1.7 fold) change the K_d_ of the CYP11B2 – Adx complex. Compared to 11-deoxycorticosterone and 11-deoxycortisol, intermediate corticosterone decreased K_d_, without affecting k_on_; and it also stabilized the complex (τ_1/2_ increased 3-4.5 times) (Table 2).

The effect of alternative substrate progesterone [24] was similar to that of 11-deoxycorticosterone on K_d_ in the case of CYP11B1 – Adx complex. Progesterone did not induce any significant changes in the CYP11B2 – Adx complex but decreased its stability (Table 2). Thus, steroid substrates similarly affect CYP11B – Adx interactions, predominantly reducing their stability (τ_1/2_) due to increase in k_off_, as shown in SPR sensorgrams (Fig. 2.5, 2.6, 2.8, 2.9).

Taking together we were able to detect different effects of steroid substrates on CYP11A1 – Adx and CYP11B – Adx interactions (Fig. 3). The substrate cholesterol accelerated the assembly of CYP11A1 – Adx complex and increased the binding affinity. In contrast, the presence of substrates (11-deoxycortisol, 11-deoxycorticosterone) reduced the stability of the CYP11B – Adx complexes without significant effect on their affinity.

**Fig. 3.**
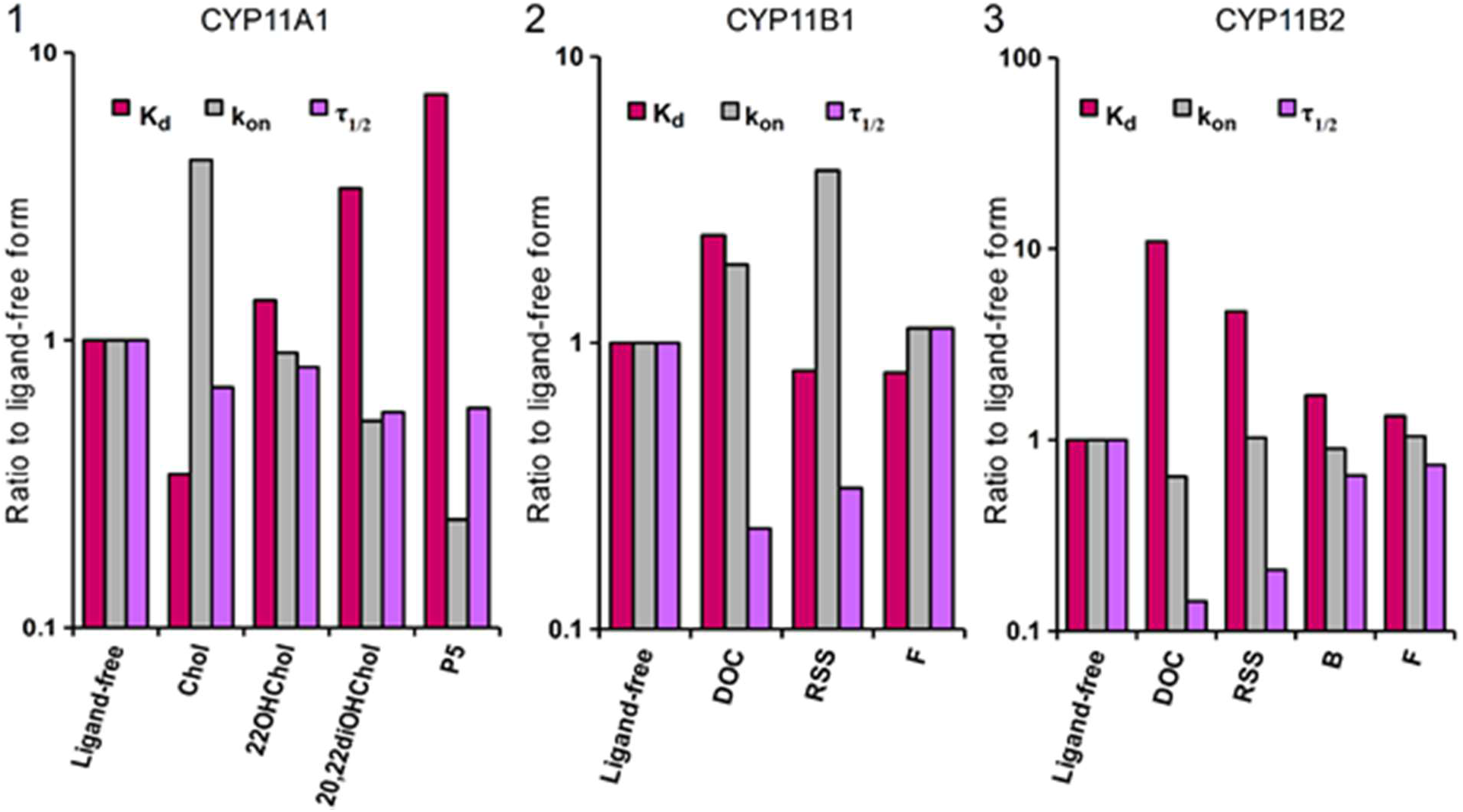
Family-specific influence of steroids on CYP – Adx interaction. 1 – CYP11A1 – Adx; 2 – CYP11B1 – Adx; 3 – CYP11B2 – Adx. The Y axis represents the ratio of the values of K_d_, k_on_, τ_1/2_ of the CYP – Adx complexes in the presence of steroids to the values of these parameters obtained for ligand-free CYP. Chol – cholesterol, 22OHChol – 22R-hydroxycholesterol, 20,22diOHChol – 20R,22R-dihydroxycholesterol, P5 – pregnenolone, DOC – 11-deoxycorticosterone, RSS — 11-deoxycortisol, B – corticosterone, F – cortisol. **Double column**

## 4. Discussion

Protein–protein interaction (PPI) inhibitors are a rapidly expanding class of therapeutics. Many essential cellular functions rely on the precise and timely interaction of proteins forming stable or transient complexes. Modulation of the PPI is therefore of fundamental importance for receptors, transcription factors, transport proteins etc. [30–34]. Among electron transfer proteins, of particular interest is the PPI involved in steroid hormones biosynthesis as many CYP proteins in this pathway are drug targets.

In the present work we demonstrated that steroid compounds modulate the interactions between three steroidogenic mitochondrial CYPs and their common redox partner Adx. The human adrenal cortex consists of three zones: the zona glomerulosa, the zona fasciculata, and the zona reticularis, which secrete mineralocorticoids, glucocorticoids, and adrenal androgens, respectively. CYP11A1 is expressed in all three zones, while CYP11B1 is present in the zona fasciculata/reticularis and CYP11B2 in the zona glomerulosa. The ratio of redox proteins in the mitochondria of the adrenal cortex was reported as AdR: Adx: CYP equal to 1: 3: 8 [20], another study estimated higher content of Adx [21]. Even with these estimations competition for electrons could not be excluded in the zona glomerulosa of the adrenal glands, where CYP11A1 and CYP11B2 are coexpressed, and in the zona fasciculata/reticularis, where CYP11A1 and CYP11B1 coexpressed [4,35]. Here we detected isoform-specific differences in the effect of substrate: steroids differently affect complex formation of Adx with CYP11A1 and CYP11B, modulating either stability or rate of complex formation. Cholesterol increased CYP11A1 affinity to Adx, while 11-deoxycorticosterone decreased CYP11B2 affinity to Adx. This is in a good accordance with the sequence of the reactions: pregnenolone must be formed first and serves as a precursor of all steroid hormones. It is also in an agreement with the previously published data, where *in vitro* enzymatic activity of human CYP11B1 and CYP11B2 decreased when CYP11B isoforms are coexpressed with CYP11A1 [36]. It is generally accepted that electrostatic interactions dominate in the interaction of Adx with mitochondrial CYPs, which is supported by the crystal structure of the complex with CYP11A1 [22]. The main difference between CYP11A1 and CYP11B structures is the conformation of the meander region at the interface with Adx highlighting its importance for isoform-specific interaction with redox partner (Fig 4A).

**Fig. 4.**
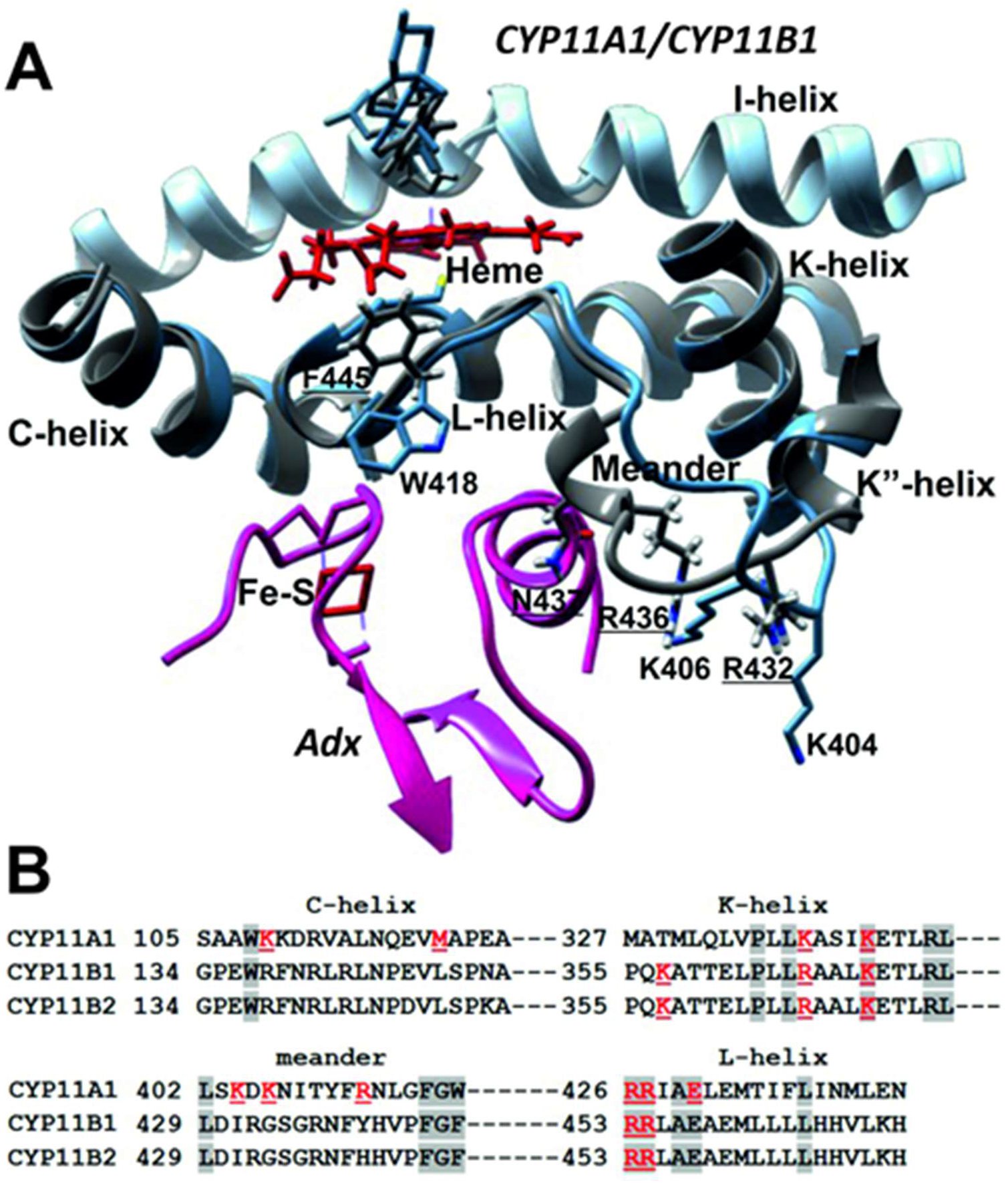
**A.** Superimposed regions of CYP11A1 (blue) (PDB ID: 3N9Y) and CYP11B1 (grey) (PDB ID: 6M7X) involved in the interaction with Adx (pink). The names of the interacting proteins are indicated in italics. Meander region represents noticeable differences between CYP11A1 and CYP11B1 (labeled as underlined text) in the amino acids involved in the complex formation with redox partner. **B.** Sequence alignment of human mitochondrial CYPs. The amino acids involved in the interaction with Adx are underlined. **1.5 column**

We suggested that different properties of intermediates with hydroxylated moiety close enough to the heme and their different modes of binding to the heme iron might affect the iron-sulfur bond with invariant cysteine residue of CYP11 enzymes and could be further translated to the proximal surface, more likely to meander, where Adx binds. This was not observed in a different experimental setup, when the effect of Adx was estimated on the heme active site but only with cholesterol by Resonance Raman spectroscopy [19].

Within CYP11 family, only CYP11B1 possesses exclusively monooxygenase function, while CYP11A1 and CYP11B2 catalyze multistep reactions (Fig. 1) [22,24,37]. Binding of cholesterol, which has rather low affinity to CYP11A1 (compared to intermediates) [38,39] enhances affinity of CYP11A1 to Adx. On the contrast, intermediates 22R-hydroxycholesterol and 20R,22R-dihydroxycholesterol bind to CYP11A1 more tightly and metabolized more efficiently [39], but being bound to CYP11A1 they decrease the affinity to Adx, supporting the idea that productive electron supply is crucial for the first and slowest step in the pregnenolone biosynthesis (Fig. 1). The intermediate 22R-hydroxycholesterol without proceeding the reaction could be released and function as the liver X receptor ligand [40]. In the case of CYP11B2, intermediate corticosterone has lower affinity to CYP11B2 (compared to 11-deoxycorticosterone) [41], but increases affinity of CYP11B2 – Adx complex, favoring the aldosterone production. In addition, substrates of CYP11B2 significantly decrease the dissociation time of the complex with redox partner to ensure effective shuttling of Adx to transfer subsequently two electrons for catalysis.

Using the SPR method we were able to detect the effect of different steroids on protein-protein interactions of three different mitochondrial CYPs with a common redox partner, which have profound functional consequences.

## 5. Conclusions

In the present work we propose that the pregnenolone synthesis as well as the production of gluco- and mineralocorticoids is modulated by the substrate-induced PPIs in mitochondrial CYP-dependent systems. This is in contrast to the microsomal CYPs, where the CYP – cytochrome P450 – reductase complex formation is not affected by the ligands [14,28]. Moreover, the effect of the substrate is isoform specific. Based on our experimental data we suggest that substrate binding contributes more to the CYP11A1 – Adx recognition, providing priority of this enzyme for the common pool of reducing equivalents. Our findings expand the current knowledge about mitochondrial CYP-redox partner interactions and provide a basis for the targeted modulation of specific CYP activity.

## Abbreviation

K_d_: apparent dissociation constant of a protein complex,
k_on_: association rate constant,
k_off_: dissociation rate constant,
SPR: surface plasmon resonance,
Chol: cholesterol,
22OHChol: 22R-hydroxycholesterol,
20,22diOHChol: 20R,22R-dihydroxycholesterol,
25OHChol: 25-hydroxycholesterol,
20OHChol: 20S-hydroxycholesterol,
DOC: 11-deoxycorticosterone,
RSS: 11-deoxycortisol,
B: corticosterone,
18OHB: 18-hydroxycorticosterone,
F: cortisol,
P4: progesterone,
P5: pregnenolone,
PPI: protein-protein interactions,
CYP: cytochrome P450,
Adx: adrenodoxin,
AdR: adrenodoxin reductase,
TN: turnover number.

## Acknowledgments

We are thankful to Petrenko A.R. from IBMC for contributing to the SPR-experiments, to Shkel T.V., Grabovec I.P. from the Institute of Bioorganic Chemistry for the excellent support in CYPs purification and functional characterization.

## Author Contributions

Yablokov E.O., Kaluzhskiy L.A., Mezentsev Y.V. performed all biosensor experiments and calculated the parameters of protein-protein interactions. Kavaleuski A.A. performed Adx and AdR purification. Yablokov E.O., Gilep A.A., Strushkevich N.V. formulated the main idea. Sushko T.A., Yablokov E.O., wrote the original draft of the manuscript. Ershov P.V., Gilep A.A., Ivanov A.S., Strushkevich N.V designed the project, supervised the data analysis and critically revised the manuscript.

## Graphical abstract

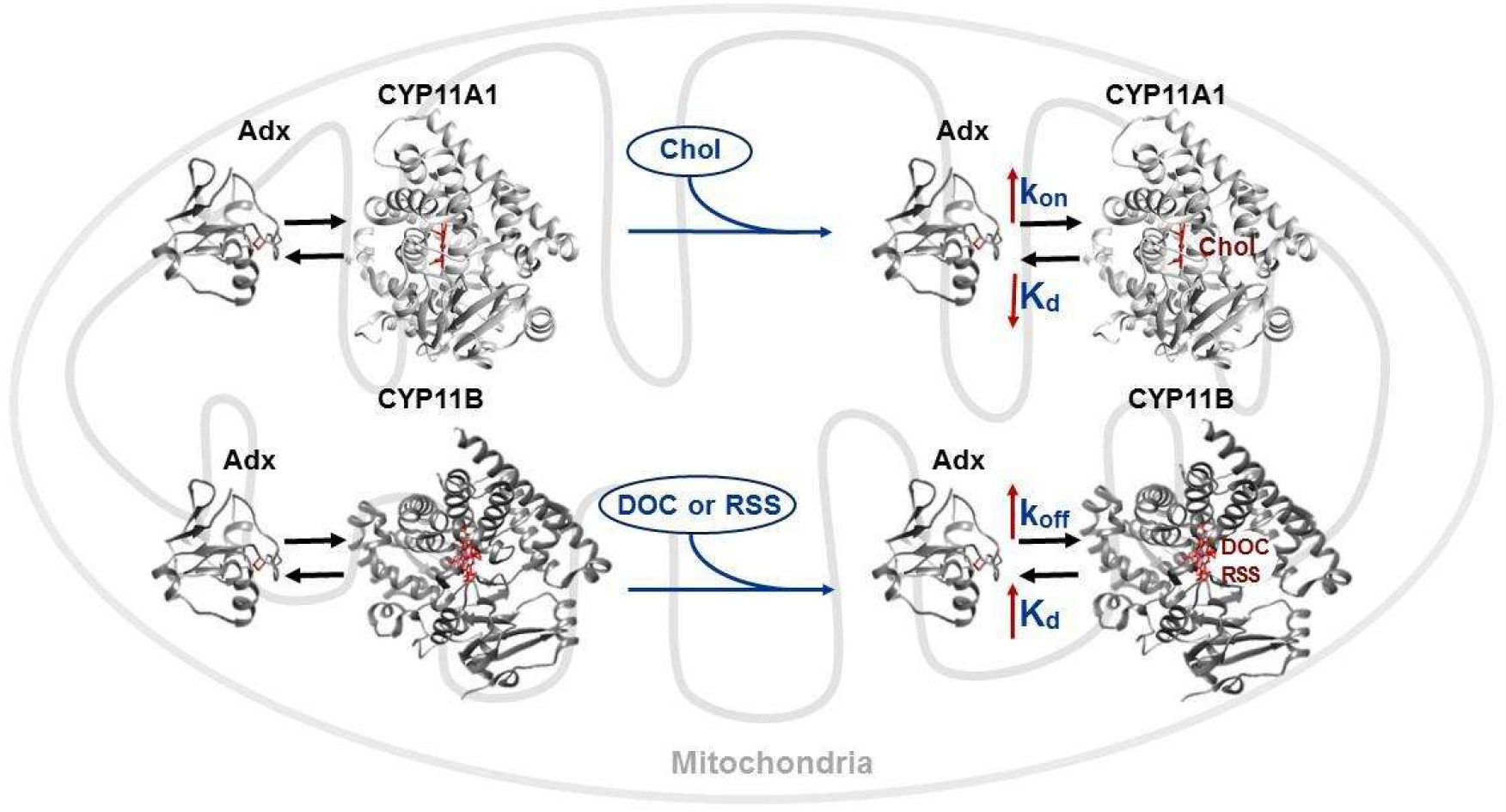

